# Temporal changes in post-alighting resistance to *Rhopalosiphum padi* (bird cherry-oat aphid) in ancestral wheats

**DOI:** 10.1101/2019.12.16.877589

**Authors:** Amma L. Simon, Lesley E. Smart, Kim E. Hammond-Kosack, Linda M. Field, Gudbjorg I. Aradottir

## Abstract

1. Aphids reduce wheat yield through feeding on phloem sap, inflicting direct and indirect damage to the plant. Currently no commercial wheat varieties have been bred for resistance to *Rhopalosiphum padi* (bird cherry-oat aphid). However, post-alighting resistance has been identified in *Triticum monococcum* lines at the seedling stage.
2. To further characterise the resistance in *T. monococcum* at plant ages, we have investigated the development, survival and reproductive success of *R. padi* on two susceptible wheats *Triticum aestivum* (variety Solstice) and *T. monococcum* MDR037, as well as on the partially resistant *T. monococcum* MDR045 and MDR049, using one, two, 12 (flag leaf) and 20 week-old (inflorescence) plants.
3. We found that the host plant resistance reduced development and reproductive success in aphids. However, the effect decreased with plant age on MDR045 but increased with plant age on MDR049.
4. The observed resistance to aphids has strong potential for introgression into commercial wheat varieties, which could have an important role in Integrated Pest Management strategies to reduce aphid populations and virus transmission.

## Introduction

Plant defence against herbivores has been classified into three different mechanisms; direct, indirect and tolerance (Boege and Marquis, 2005; Stamp, 2003; Strauss and Agrawal, 1999). These mechanisms lead to the expression of resistance and tolerance traits in plants (Agrawal, 2007; Czesak et al., 2008; Rasmann and Agrawal, 2009). Plants with pre-alighting resistance (antixenosis) use indirect defence mechanisms, involving visual cues and/or the production of Volatile Organic Compounds (VOCs) that are emitted from the plant, and affect insect behaviour, for example, causing the herbivores to express non-host behavioural responses (Kogan and Ortman, 1978). Plants with post-alighting resistance (antibiosis) to insects use direct defence mechanisms, either reducing the ability of the insect to access the food source, reducing the quality or digestibility of the food, or making the food source toxic by the production of phytochemicals. This leads to a decrease in insect development and/or reproduction (Painter, 1951). Whilst pre- and post-alighting resistance can prevent herbivory, tolerance merely reduces the adverse effects that herbivory has on a plant (Strauss and Agrawal, 1999).

In response to herbivory, plants have developed various defence mechanisms which lead to expression of resistance and tolerance traits in plants. Both the plant defence theory and plant–herbivore coevolutionary theory assume that resistance traits have evolved as adaptations to reduce herbivory (Czesak et al., 2008), which has been evidenced in studies showing that resistance traits are under selection from herbivores (Agrawal, 2007; Rasmann et al., 2015; Thaler, 1999; Turley and Johnson, 2015; Züst and Agrawal, 2016). Such co-evolution has created an evolutionary arms race between aphids and their hosts (Czesak et al., 2008) leading to rapid diversification in both Angiosperms and aphids during the early Cretaceous period (Dixon, 1998).

Optimal defence theory (ODT) predicts the allocation of chemical defence against insects, within the context of plant fitness. It is assumed that changes in resource allocation will be determined by the rate of herbivorous attack in the absence of defence (Boege and Marquis, 2005; Tiffin, 2002), the importance of the organ under attack for development and reproduction (Boege and Marquis, 2005) and the cost of defence allocation (Koricheva, 2002). ODT predicts that plant age and genotype can affect resistance strategies (Boege and Marquis, 2005; Farnsworth, 2004; Koricheva, 2002; Tiffin, 2002). It has been theorised that younger plants have better anti-herbivory defence (Boege and Marquis, 2005) as seedlings need more protection (Iwasa et al., 1996; Nicole M. van Dam et al., 1995), this can also be the case for younger leaves on the same plant (van Dam et al., 1996). On the other hand, on some genotypes herbivory occurs more on seedlings than adult plants (Fenner et al., 1999), similar to the phenomenon of Adult Plant Resistance which has been observed in plant-pathogen interactions (Dyck, 1979; Dyck and Samborski, 1979). These contrasting theories emphasise that the genotype of both the herbivore and plant plays an important role in whether younger or older plants have increased anti-herbivory defence (Koricheva, 2002).

Wheat is a main ingredient in many diets worldwide and so is an important crop for food security (Shewry, 2009; Shewry and Hey, 2015). However, its production is often compromised by weed competition and attack from pathogens and insects. The most important insect pests are aphids and their control currently relies on the use of insecticides, although there are increasing reports of insecticide resistance (Foster et al., 2014). This, alongside restrictions on the use of some insecticidal classes (Pickett, 2013), could culminate in limited options for pest population control. Most of the ~4,000 extant aphid species (Blackman and Eastop, 2007) are phloem feeders and have developed methods of bypassing plant defences so that they can remain at a phloem site for days (Tagu et al., 2008). Whilst feeding, aphids can transmit viruses, for example Barley Yellow Dwarf Virus (BYDV) (Fiebig et al., 2004), which is transmitted by cereal aphids including Rhopalosiphum *padi* (L.) (Chapin et al., 2001; Halbert et al., 1992; Leather et al., 1989). Aphids also secrete honeydew which attracts saprophytic fungi that cover the leaves, reducing the plant’s photosynthetic ability (Rabbinge et al., 1981). Through a combination of damage, *R. padi* can cause up to 30% yield losses in cultivated cereals (Leather et al., 1989; Voss et al., 1997).

Commercial hexaploidy wheat has not been bred with natural resistance to *R. padi* in mind, however, studies have shown that the diploid ancestral species *Triticum monococcum* (L.) (genome A^m^ A^m^) has varying levels of post-alighting resistance to aphids (Sotherton and Lee, 1988) including *R. padi* (Greenslade et al., 2016; Radchenko, 2011). For *R. padi*, *T. monococcum* lines MDR045 and MDR049 have been shown to have partial resistance in one week old seedlings (Greenslade et al., 2016; Aradottir per comms.) However, it is not known whether this resistance is stable through plant development. It is important to know this, as according to ODT, herbivory, plant age and plant genotype can affect resistance strategies (Boege and Marquis, 2005; Farnsworth, 2004; Koricheva, 2002; Tiffin, 2002). The study reported here, explored the development, survival and reproduction of *R. padi* on two susceptible wheat *Triticum aestivum* (variety Solstice) and *T. monococcum* MDR037, and on the partially resistant *T. monococcum* MDR045 and MDR049 at four different stages of plant development, testing the hypothesis that the post-alighting resistance is subject to temporal control.

## Materials and methods

### Plants and insects

Cultures of non-viruliferous adult *R. padi* were reared on *Hordeum vulgare* (L.) variety Saffron. *Triticum monococcum* MDR037, MDR045 and MDR049 were provided by the Wheat Genetic Improvement Network (WGIN) and *T. aestivum* var. Solstice was provided by Rothamsted Research, all seeds were stored at 4°C.

Plants used for testing at one and two weeks post sowing were grown in 2cm diameter seed trays in a controlled environment room at 20°C (± 1°C),16:8 h light: dark photoperiod with 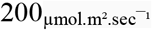 light intensity and watered with approximately 8ml daily. Plants for testing at later growth stages were grown in 2cm seed trays in the controlled environment room for seven days, before being vernalised for six weeks. After vernalisation, plants were transferred to 12.5cm diameter pots and grown in a glass house until the start of experimentation when they were moved back to the controlled environment room with the same environmental conditions as before. Experiments on different plant ages were done independently and sequentially due to limited controlled environment space. The growing medium was 75% medium grade peat, 12% screened sterilised loam, 3% medium grade vermiculite and 10% grit with an N content of 14% and a P_2_O_5_ content of 16% (from Petersfield Products, Cosby, Leicester).

### Assessment of aphid development, survival and reproductive success

Development, survival and reproductive success bioassays were carried out in a controlled environment room (20°C (± 1°C), 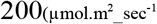 for 16:8h L:D) on all four wheat genotypes at the four different ages; one week, two week, 12 week and 20 week. Experiments on each growth stage was done independently, in sequence due to limited space, however all experiments were carried out in the same controlled environment room. For each experiment, ten plants (biological replicates) of each wheat genotype were placed in a randomised experimental design.

Three to five mature alate aphids were placed on each plant in a 2cm clip cage and allowed to produce nymphs overnight. The following morning the mature aphids were removed, and the all neonate nymphs (<1 day old) were counted and removed carefully from plants with a fine-haired paintbrush, taking care not to damage them or their stylets, placed in pre-weighed Eppendorf tubes and weighed using a Microbalance (Cahn 33; Scientific and Medical Products Ltd, Manchester, UK) to determine the average nymph weight.

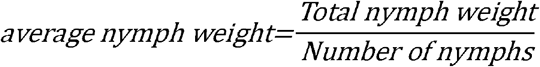

After weighing, nymphs were transferred back to their respective plants using a fine-haired paintbrush and left undisturbed for six days. After six days, the number of survivors was recorded, and they were re-weighed to determine the mean Relative Growth Rate (mRGR). This was calculated (Radford, 1967; Leather & Dixon, 1984) as:

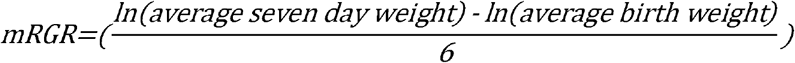

The % survival after six days was calculated as:

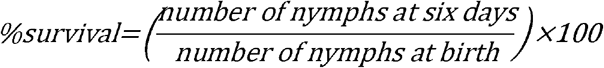

After weighing, one of the nymphs, chosen at random, was transferred back to their original plant, left undisturbed to develop, and monitored each day. The time taken to produce their first nymph and the number of nymphs they produced over their lifetime were recorded. Nymphs were removed to prevent overcrowding. All aphids developed into adult rapturous aphids.

### Statistical analysis

The differences in aphid mRGR, the percentage survival after six days, the number of days to produce a first nymph and the number of nymphs produced were examined using a two-factor analysis of variance (ANOVA) with Tukey’s HSD post hoc in which the plant age and the plant genotype were factors, interactions between factors was also included in analysis. Data were subjected to transformation to comply with assumptions of normality and equal variance. The number of days to produce the first nymph was log transformed, the number of nymphs was square root transformed and percentage survival logit transformation before statistical analysis. Confidence level of P<0.05 was considered to be statistically significant. GenStat® (2016, 18^th^ Edition, © VSN International Ltd, Hemel Hempstead, UK) statistical software was used for analysis.

## Results

### Aphid development

Overall, plant age alone had no effect on mean relative growth rate (mRGR), however wheat genotype affected mRGR, as aphids feeding on MDR049 had lower mRGR than on Solstice (*F*_3,134_ = 5.13, *P* < 0.05) (Fig. 1).

**Figure 1.**
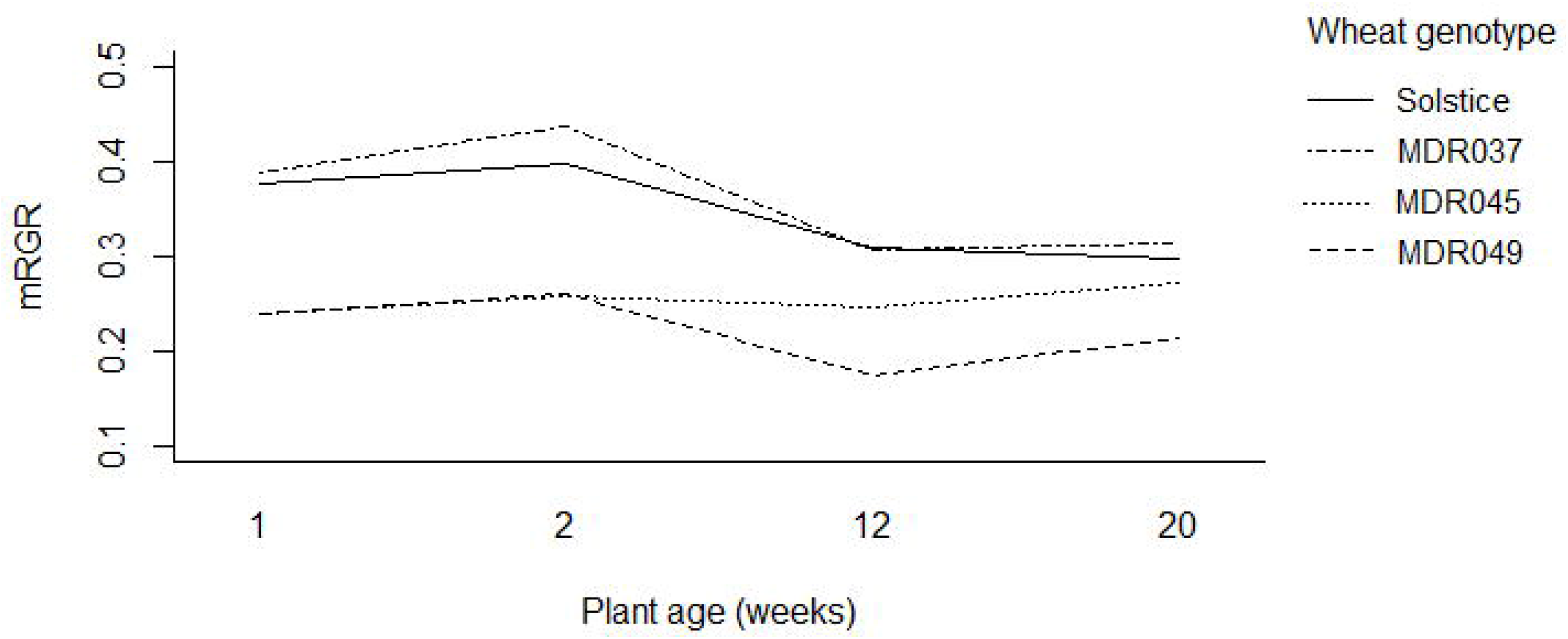
Mean Relative Growth Rate (mRGR) of *Rhopalosiphum padi* on one week old, two week old, 12 week old and 20 week old *Triticum monococcum* MDR037, MDR045 and MDR049 and *Triticum aestivum* Solstice at four growth stages. Error bars removed for clarity.

There was an interaction between wheat age and genotype for mRGR (*F*_9,134_ = 0.75, *P* < 0.05) (Fig. 1). Aphids on one and two-week old MDR045 had lower mRGR than on Solstice (*F*_3,134_ = 7.79, *P* < 0.05) (Fig. 1).

### Aphid survival

Wheat genotype had no effect on aphid survival, however, plant age affected the survival (*F*_3,134_ = 2.67, *P* < 0.05). There was also an interaction for how well aphids fared on the plants (*F*_9,134_ = 1.98, *P* < 0.05) (Fig. 2). Plant age only affected the survival of aphids feeding on MDR049, which was higher on two-week old plants than on 12-week old plants (*F*_3,134_ = 2.78, *P* < 0.05) (Fig. 2). Aphids on 12 week-old MDR045 and those on 20 week-old MDR049 had lower survival than those on 12 and 20 week-old Solstice (*F*_3,134_ = 0.71, *P* < 0.05) (Fig. 2).

**Figure 2.**
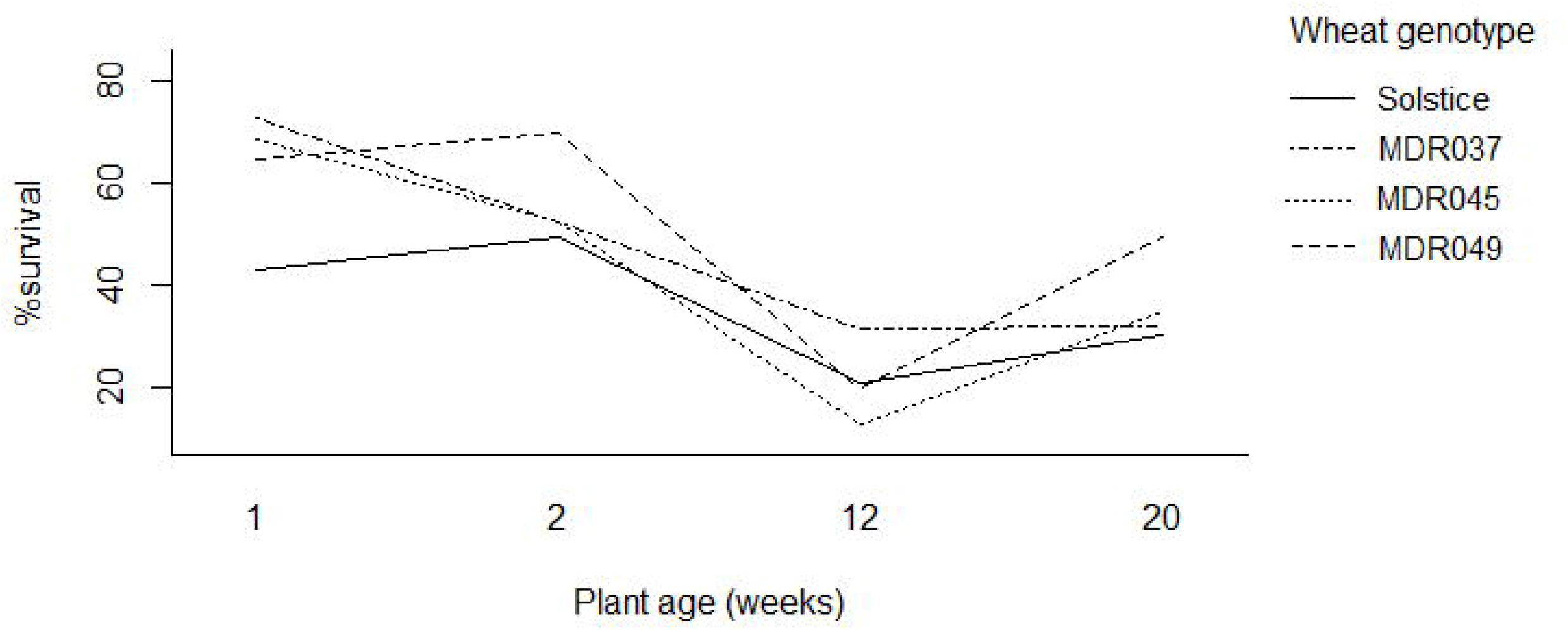
Mean percentage survival of *Rhopalosiphum padi* on one week old, two week old, 12 week old and 20 week old *Triticum monococcum* MDR037, MDR045 and MDR049 and *Triticum aestivum* Solstice at four growth stages. Error bars removed for clarity.

### Aphid reproductive success

Aphids on older (12 and/or 20 weeks) plants took longer to produce their first nymph (*F*_3,121_ = 38.34, *P* < 0.001) (Fig. 3) and produced fewer nymphs (*F*_3,121_ = 16.62, *P* < 0.001) (Fig. 4) than those on younger (one and/or two-weeks) plants. Wheat genotype had no effect of number of days it took aphids to produce the first nymph, however, aphids on MDR049 produced fewer nymphs than aphids on MDR037 and Solstice (*F*_3,121_ = 14.17, *P* < 0.01) (Fig. 4)

**Figure 3.**
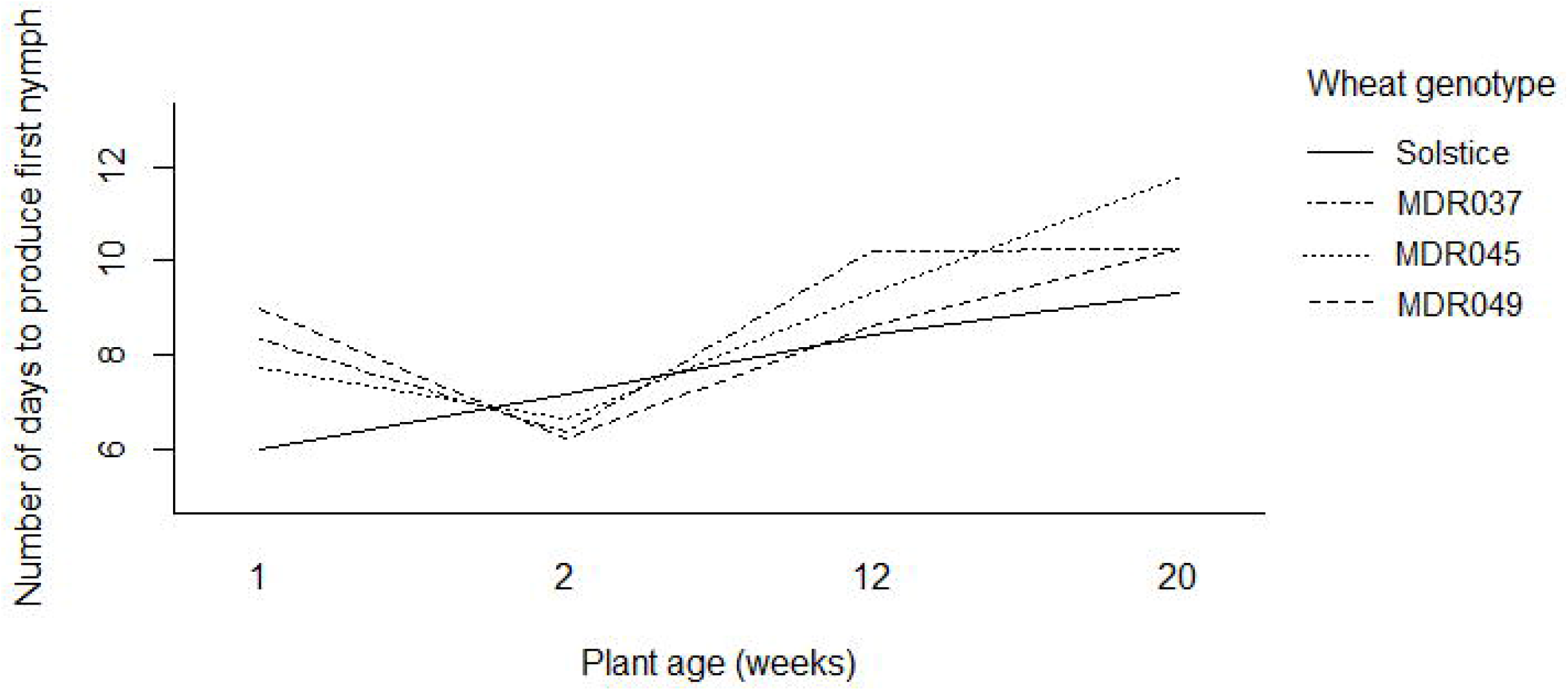
Mean number of days *R. padi* took to produce their first nymph on one week old, two week old, 12 week old and 20 week old *Triticum monococcum* MDR037, MDR045 and MDR049 and *Triticum aestivum* Solstice at four growth stages. Error bars removed for clarity.

**Figure 4.**
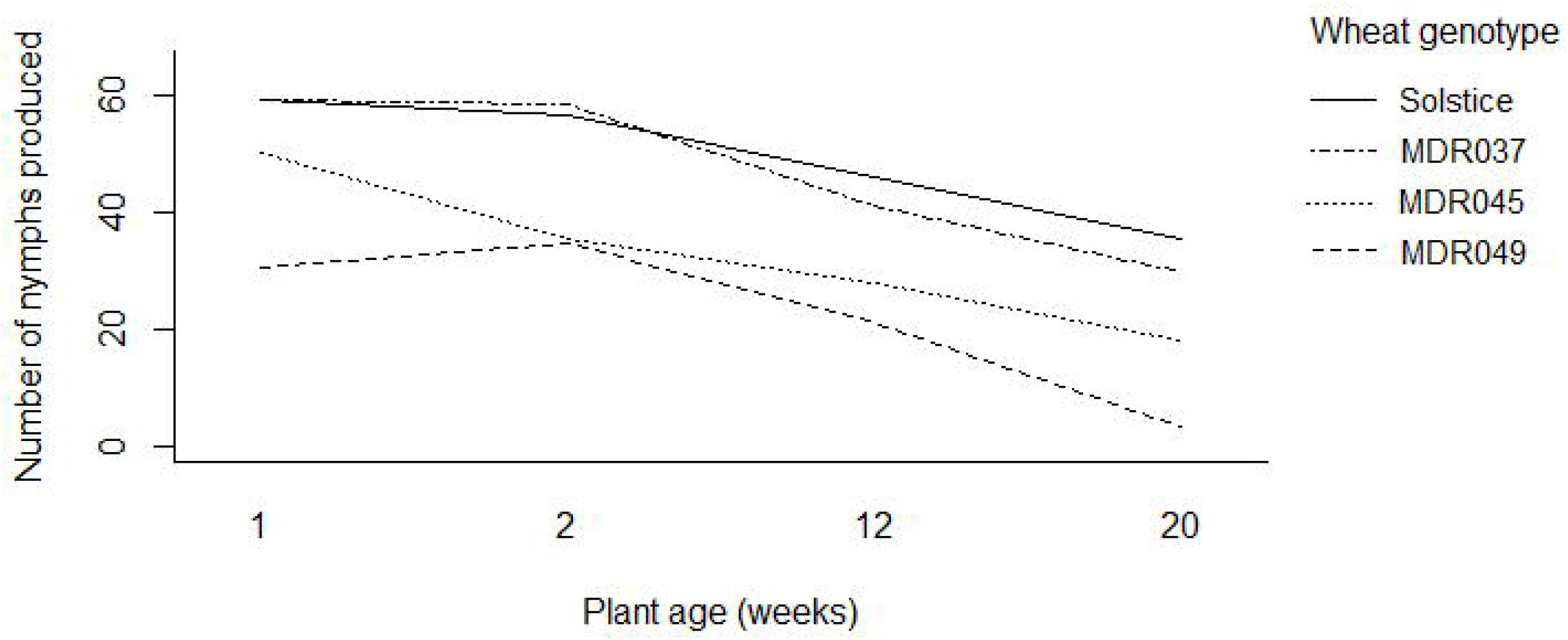
Number of nymphs produced by *Rhopalosiphum padi* on one week old, two week old, 12 week old and 20 week old *Triticum monococcum* MDR037, MDR045 and MDR049 and *Triticum aestivum* Solstice at four growth stages. Error bars removed for clarity.

There was an interaction between the age and wheat genotype for time taken to produce the first nymph (*F*_9,121_ = 2.37, *P* < 0.05) (Fig. 3). Aphids on one week-old Solstice produced their first nymph quicker than those on MDR037, MDR045 and MDR049 (*F*_3,121_ = 2.46, *P* < 0.01) (Fig. 3) and aphids on 20 week-old MDR045 took longer to produce their first nymphs than those on Solstice (*F*_3,121_ = 2.46, *P* < 0.05) (Fig. 3).

There was also an interaction between the age and wheat genotype for nymph production (*F*_9,121_ =1.04, *P* < 0.05) (Fig. 4). Aphids on MDR045 produced fewer nymphs than those on Solstice and MDR037 (*F*_3,121_ = 14.17, *P* < 0.05) (Fig. 4) on one, two and 12 week-old plants with significant differences decreasing with plant age. On the other hand, aphids on MDR049 produced fewer nymphs than those on Solstice and MDR037 at all growth stages (*F*_3,121_ = 14.17, *P* < 0.01) (Fig. 4) with increased significance on 12 and 20 week-old plants

## Discussion

Screening for resistance to aphids in *T. monococcum* is often only done at a single growth stage (di Pietro et al., 1998; Greenslade et al., 2016; Migui and Lamb, 2003; Simon et al., 2017), but it is also important to understand the interaction between plant age and resistance, hence, here we report on how plant age affects *R. padi* development and reproductive success on ancestral *T. monococcum* plants at four different growth stages for four different wheat genotypes. Our main finding was that this can depend on plant genotype as well as plant age. We show that for *T. monococcum* MDR045, aphid resistance was stronger in seedlings, however, in *T. monococcum* MDR049 aphid resistance increased with plant age and was stronger in plants at the inflorescence stage.

The Optimal Defence Theory (ODT) assumes that defence is costly (McCall and Fordyce, 2010) and so allocation to defence will occur when there is a low cost: benefit ratio (McCall and Fordyce, 2010; Stamp, 2003). The more important a tissue is for the fitness of the plant, the higher the concentrations of defence compounds are likely to be (Boege and Marquis, 2005). It has been suggested that younger leaves have higher defences than older leaves (Nicole M van Dam et al., 1995) because of their importance to the plant’s establishment (Iwasa et al., 1996) and its increased photosynthetic ability (Wiedemuth et al., 2005). This contradicts our findings, as across all genotypes, *R. padi* on younger wheat plants (1 week and 2 week-old plants) produced their first nymph quicker and produced more nymphs than those on older plants. This is also in contradiction to previous work by Leather and Dixon (1981) which showed that *R. padi* was less fecund on wheat seedlings at GS11 than those on older plants, however, in these experiments *Tritium aestivum* cultivar Maris Huntsman was used. This emphasises the importance of cultivar and genotype effects on aphid life history traits. The higher *R. padi* fecundity on seedlings observed may be because *R. padi* is associated with feeding on cereal crop seedlings during warm autumns leading to BYDV transmission (Fabre et al., 2003; Riedell et al., 1997) and could be better adapted to feeding on seedlings than on older plants.

It has previously been shown that *T. monococcum* seedlings have resistance to *R. padi* (Greenslade et al., 2016) and *S. avenae* (di Pietro et al., 1998; Migui and Lamb, 2004; Simon et al., 2017), however, post-alighting resistance in adult *T. monococcum* has only been observed towards *S. avenae* (Migui and Lamb, 2004, 2003). Here we have shown that *T. monococcum* MDR049 had reduced mean relative growth rate (mRGR) and produced fewer nymphs than the susceptible Solstice and/or *T. monococcum* MDR037 at all plant ages indicating that this resistance is stable in MDR049. This contradicts Migui and Lamb, (2003) study, where no resistance to *R. padi* was observed in adult *T. monococcum*, but in that study they only considered aphid biomass. This emphasises the need for in depth investigations of both aphid development and reproductive success, for each genotype, when exploring the interaction between plants and aphids with the aim to identify resistance.

Antibiosis in MDR049 seedlings has been studied using an electrical penetration graphs and linked with the inability for aphids to carry out sustained feeding (Greenslade et al., 2016; Simon et al., 2017), and smaller vascular bundles (Simon et al., 2017), however, it remains to be tested whether this effect on aphid feeding behaviour is consistent at all growth stages.

Whilst in some plant species, it has been shown that plants become more susceptible to herbivores with age (Iwasa et al., 1996; McCall and Fordyce, 2010), with higher levels of defence compounds found in seedlings of Ribwort plantain (*Plantago lanceolata* (L.)) (Deane Bowers and Stamp, 1993), gypsy flower (*Cynoglossum officinale* (L.)) (Nicole M. van Dam et al., 1995) and diesel tree (*Copaifera langsdorfii*(D.)) (Macedo and Langenheim, 1989) and decreased herbivory by *Stenoma assignata* (M.) larvae on *C. langsdorfii* (Macedo and Langenheim, 1989), generalist slug (*Deroceras reticulatum* (M.)) on Prostrate knotweed (*Polygonum aviculare* (L.)), acyanogeic white clover (*Trifolium repens* (L.)) and lettuce (*Lactuca sativa* (L.)) (Fenner et al., 1999). On the other hand, some adult plants have been shown to have higher defence compounds (Boege, 2005) and herbivores prefer to feed on and do better on seedlings rather than on older plants including *D. reticulatum* on Corn daisy (*Glebionis segeutum* (L.)), Stinking willie (*Jacobaea vulgaris* (L.)), Red clover (*Trifolium pratense* (L.)), Chickweed (*Stellaria media* (L.)), Catgrass (*Dactylis glomerata* (L.)) and Poppy (*Papaver rhoeas* (L.) (Fenner et al., 1999). Geometridae chewing insects on the tree species *Casearia nitida* (Boege, 2005) and the Dusky Arion slug (*Arion subfuscus*) on willow also performed better on adult plants (Fritz et al., 2001). These examples show that defence allocation can be complex and dependent on age, herbivory and plant genotype (Boege and Marquis, 2005; Farnsworth, 2004; Koricheva, 2002; Tiffin, 2002). Here we show a combination of these within the same cereal species, where there was an interaction between wheat genotype and age for all parameters. The interaction is most apparent when looking at nymph production on MDR045 and MDR049 over the four growth stages. We have shown that antibiosis resistance affecting *R. padi* nymph production in MDR045 decreased with age, whereas on MDR049, it increased with plant age, with increased significance between susceptible lines and MDR049 in 12 and 20 week-old plants. Similarly, di Pietro *et al.*, (1998) showed that *T. monococcum* TM44 and TM46 seedlings had antibiosis to *S. avenae* but as adult plants, there was only a reduction in development for aphids feeding on TM46 (Migui and Lamb, 2003).

The increased resistance with age observed with MDR049 may be due to a form of Adult Plant Resistance (APR) (Dyck, 1979; Dyck and Samborski, 1979), as reported in response to pathogen infections. For example, the APR gene *Lr34/Yr18* in *T. aestivum* PI250413 confers resistance to stem rust (Dyck and Samborski, 1979) and in *T. aestivum* RL6058, RL6077 and Line902 confers resistance to stripe rust (Singh, 1992). The effect of the APR gene alone can vary depending on the level of infection (Ellis et al., 2014; Singh et al., 2014) but combining various APR genes can cause high levels of resistance to rust (Singh et al., 2014). Single APR genes can interact with major resistance (*R*) gene mediated defences, increasing the phenotype associated with *R* genes and resulting in lower infection of the pathogen (Ellis et al., 2014; Singh et al., 2014). Little is known about APR towards insects but in our work, MDR049 showed resistance throughout all growth stages which may be due to the presence of *R* gene(s), which provide resistance from seedling to adult plant, but in the adult plant the resistance was increased by the expression of APR gene(s).

The presence and level of resistance in *T. monococcum* MDR045 and MDR049 is dependent on the plant growth stage and the plant genotype. Resistance that reduces nymph production decreased with age in MDR045 however, this resistance increased in later growth stages in MDR049. This highlights the need to consider the variability of genotype responses over different growth stages as well as examining both aphid development and reproductive success when investigating resistance to insect pest species.

## Author Contributions

A.L.S and G.I.A designed the experiments. A.L.S conducted experiments, analysed data and wrote the manuscript. L.S, K.H-K, L.F and G.I.A supervised the project and provided critical corrections of the manuscript.

## Acknowledgements

We would like to thank Janet Martin for assistance in experimental design, Suzanne Clark for statistical advice. Jill Maple, Jack Turner, Steve Harvey, Fiona Gilzean, Irene Castellan and Emmanuel Ziramba for assistance in plant care during vernalisation and growing plants to the required growth stage. John Foulkes for providing critical corrections of the manuscript. The *T. monococcum* resources were developed within the core project of the Wheat Genetic Improvement Network (http://www.WGIN.org.uk), which is supported by a grant from the Department for Environment, Food and Rural Affairs (Defra, AR0709) and is based on phenotyping data developed as part of the Designing Future Wheat Institute Strategic Programme (BBS/E/C/000I0250) funded by the UK Biotechnology and Biological Sciences Research Council (BBSRC). This project was funded by the Lawes Agricultural Trust, University of Nottingham and the BBSRC as part of a University of Nottingham Doctoral Training Partnership PhD.

